# Age-related complexity of the resting state MEG signals: a multiscale entropy analysis

**DOI:** 10.1101/2022.09.14.507986

**Authors:** Armin Makani, Amir Akhavan, Farhad Shahbazi, Mohammad Noruzi, Marzieh Zare

## Abstract

The effects of aging on the brain can be studied by examining the changes in complexity of brain signals and fluid cognitive abilities. This paper is a relatively large-scale study in which the complexity of the resting-state MEG (rsMEG) signal was investigated in 602 healthy participants (298 females and 304 males) aged 18 to 87. In order to quantify the brain signals’ complexity, the multiscale entropy is applied. This study investigates the relationship between age and fluid intelligence with brain complexity and the variations of the complexity asymmetry between the left-right brain hemispheres across the life span. In the analysis of the brain signals, the gender difference was considered. The results showed that the complexity of rsMEG decreases across the lifespan. However, the complexity difference between the left-right brain hemispheres positively correlates with age. Furthermore, the results demonstrated that fluid intelligence and age have a positive correlation. Finally, the frequency analysis revealed a significant increase in the relative power of low and high gamma rhythms in females compared to males in all age groups.

## 1. Introduction

Various factors lead to tremendous structural changes in the human brain. Aging, unhealthy lifestyles such as alcohol consumption, neurodegenerative and infectious diseases are among the reasons for these changes [1-3]. In response to these structural changes, the function of the brain change[4]. Researchers use different medical techniques depending on whether they want to study structural or functional changes. Electroencephalography (EEG) and magnetoencephalography (MEG) are more suitable than other methods for investigating the functional changes in the human brain due to higher temporal resolution, non-invasiveness and direct measurement of neural activity [5-8].

One of the ways to study the effect of aging on the brain is to examine changes in the complexity of brain signals across the lifespan. Researchers use various linear and nonlinear measures to quantify the complexity of brain signals. Since brain signals are nonlinear and chaotic, linear measures alone cannot characterize their dynamics. As a result, using nonlinear measures has attracted growing attention in brain signal analysis. Filippo Zappasodi et al. [8] used the Higuchi fractal dimension to measure the complexity of the EEG signal of eyes-open resting-state healthy individuals. They showed that the relationship between age and EEG complexity is a parabolic function. Hideyuki Hoshi et al. [9] examined the relationship between age and source power of the resting-state MEG signal in both opened and closed eyes. They observed that the power of faster oscillations increased in the rostral area and decreased in the caudal area during aging. Yanbing Jia et al. [10] used sample entropy to examine the complexity of the resting-state fMRI signal. They showed that, unlike healthy subjects, the complexity in amygdala-cortical connectivity in patients with schizophrenia was not negatively correlated with age.

Researchers have proposed various entropy-based measures for the nonlinear analysis of brain signals[11]. Pincus [12] suggested approximate entropy to measure the complexity of dynamical systems. Then Richman et al. [13] proposed sample entropy, an improved version of approximate entropy that does not consider self-matchings. These two measures were suitable for analyzing short and noisy signals compared to other measures. However, the problem was that they were only suitable for examining short-range temporal dynamics. Finally, Costa et al. [14] introduced multiscale entropy(MSE), allowing long-range temporal dynamics to be evaluated. This measure provides information about the predictability of time series over different temporal scales.

Multiscale entropy has been applied successfully in numerous brain signal analysis studies. Tetsuya Takahashi et al. [15] investigated the EEG signal complexity of individuals affected by photic stimulation using MSE. They showed that complexity decreases with age. Catarino et al. [16] used MSE of the EEG signal to discriminate healthy individuals and patients with autism. Vladimir Miskovic et al. [17] showed that changes in the complexity of brain signals throughout the sleep cycle depend on the time scale. Chao Gu et al. [18] examined the complexity of the EEG signal of patients with ADHD and healthy participants. They found that in the resting state, the EEG complexity of healthy subjects is higher than that of patients with ADHD. In contrast, the opposite occurs during activity.

Here, we study the largest neuroimaging dataset recorded by the Cambridge Center for Aging and Neuroscience (Cam-CAN). The main advantage of this dataset is the large number of subjects in the study and the inclusion of different neural recording and imaging modalities [19]. We primarily aimed to characterize the complexity of rsMEG signals using MSE and examine its relationship with aging, changes across hemispheres, and relative power across genders. Since researchers have shown that aging leads to fluid intelligence alterations, we finally explored complexity’s relationship with fluid intelligence, which, to the be best of our knowledge, is an overlooked topic. The paper is organized as follows. In section 2, we introduce the materials and methods. Section 3 presents the results, section 4 is devoted to the discussion.

## 2. Materials and methods

### 2.1. Data and participants

All MEG datasets were collected at a single site (MRC-CBSU) using a 306-channel VectorView MEG system (Elekta Neuromag, Helsinki), consisting of 102 magnetometers and 204 orthogonal planar gradiometers, located in a light magnetically shielded room (MSR). During the resting state recording, participants sat on a chair with their eyes closed for approximately 8 min and 40 sec. Data were sampled at 1000 Hz and a band-pass filter with cutoff frequency of 0.03-330 Hz was applied. We considered a sensor array of 102 pairs of planar gradiometers. Notch frequencies associated with the alternating current of the electricity network were 50 Hz. The number of participants was 623, whose ages ranged from 18 to 88 and the number of participants in each group is presented in Table 1 [19]. Computations were carried out using the Python programming language (Python Software Foundation. Python Language Reference, version 3.7.2. available at https://www.python.org/). Besides rsMEG signals, fluid intelligence data was also used in this study. Fluid intelligence was measured by the Culture Fair Intelligence Test (CFIT) [20].

**Table 1:** number of participants in each age group.

| Group Name | Age Range | Males’ Number | Females’ Number |
| --- | --- | --- | --- |
| **G1** | 18-27 | 20 | 23 |
| **G2** | 28-37 | 48 | 48 |
| **G3** | 38-47 | 49 | 47 |
| **G4** | 48-57 | 48 | 47 |
| **G5** | 58-67 | 46 | 48 |
| **G6** | 68-77 | 48 | 47 |
| **G7** | 78-87 | 45 | 38 |

### 2.2. Preprocessing

In this study, only magnetometer channels were used to reduce the computational cost [21, 22]. In order to preprocess the data for each magnetometer sensor, we performed zero-phase filtering by applying a fifth-order band-pass Butterworth filter in both the forward and reverse directions. The cut-off frequency was between 0.5 to 90 Hz. Then the signals were decimated by a factor of 3.

### 2.3. Multiscale Entropy

For each magnetometer sensor, multiscale entropy was calculated using the proposed algorithm by costa [14]. In order to calculate multiscale entropy, three steps must be taken, including coarse-graining of the original time-series, calculation of the sample entropy for all coarse-grained signals, and summation of sample entropies for different scales. These three steps are detailed below:

a) Coarse-graining

Consider the time series *x*(1), *x*(2), …, *x*(*N*) of length *N*. The coarse-grained time series in scale *τ* 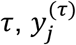,is defined as follows:

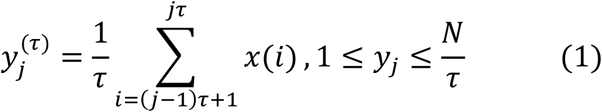

Fig. 1, A and B illustrate the coarse-graining for scales 2 and 3, respectively.

**Figure 1.**
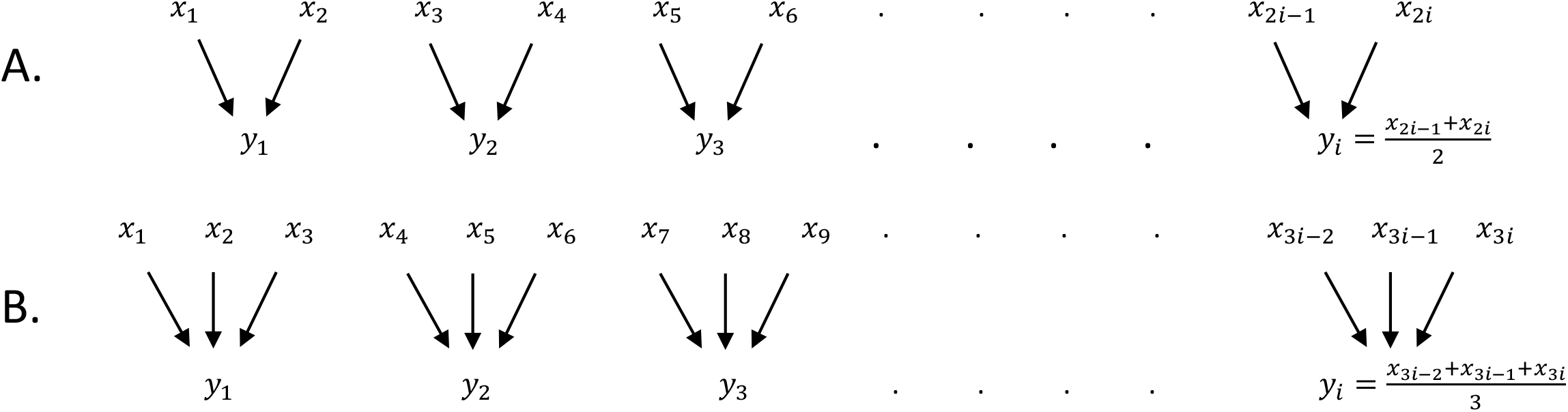
Coarse graining procedure. “x” is the original time series (coarse-grained for scale factor 1), and “y” is the new time series coarse-grained with (A) scale 2 and (B) scale 3.

b) Sample Entropy

In order to compute the sample entropy of a time series, *x* = {*x*_1_, *x*_2_, …, *x*(*N*)}, with length *N*, the first step is to embed the signal into the phase space with vectors *x*_*m*_(*i*) = [*x*(*i*), *x*(*i* + 1), …, *x*(*i* +*m* − 1)] in which m is the embedding dimension. Then the sample entropy is computed as follows:

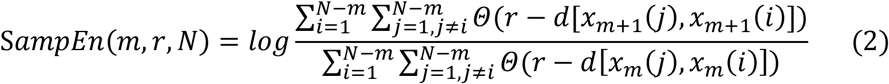

Where Θ, *d* and *r* are the Heaviside step function, the Chebyshev distance of the two vectors, and the similarity criterion, respectively. The Chebyshev distance is a metric defined on a vector space that measures the greatest distance between two vectors along any coordinate dimension [12, 13, 23].

C) Multiscale Entropy (MSE)

In order to find the multiscale Entropy (MSE) of a time series, one has to obtain coarse-grained time series from scale 1 to the desired scale and then calculate the sample entropy of each coarse-grained time series. According to Eq. 3, the sum of the sample entropies is called the multiscale entropy, and we regard it as the complexity of the time series:

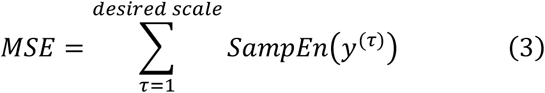

### 2.4. Relative power

The relative power was calculated in six frequency bands including delta (0.5-3Hz), theta (4-7Hz), alpha (8-13Hz), beta (15-25Hz), low gamma (30-60Hz) and high gamma (60-90Hz). The power spectral density (PSD) of different MEG rhythms was estimated using the Welch method. In this method, Hanning windows with 60% overlap were used. Also, the window length was selected as 4 seconds. In order to compute the power, the area under the PSD is computed. The relative power of each band is determined as follows:

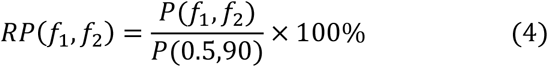

Where *P* (.) represent power, *R P* (.) stands for relative power and *f*_1_ and *f*_2_ are the low and high frequency bands respectively.

### 2.5. statistical analysis

In order to evaluate the topographic changes of MSE with age and selection of meaningful sensors, the population was divided into seven groups according to the ten-year age range (Table 1). The one-way analysis of variance was used to calculate P-values. Then Benjamini and Hochberg corrections were applied on the computed P-values to control the False Discovery Rate(FDR) [24]. In all parts of this study, sensor selection was made similarly. Meaningful sensors were used to plot the mean multiscale entropy graphs by age. In order to apply the linear regression in scatter plots and test the null hypothesis, the Python libraries Scipy and Statsmodels were used [25]. Pearson correlation coefficient was also calculated to examine the linear correlation.

## 3. Results

### 3.1. Parameter Setting

As mentioned earlier, the scale and the embedding dimension are parameters that must be determined before calculating the MSE. In order to investigate the effect of embedding dimension on MSE, one male and one female subject were randomly selected from each age group. For each participant, the MSE of all magnetometer sensors was calculated for different dimensions, with the coarse-graining factor up to 125. The average of those multiscale entropies was calculated. Since the results for different age groups were similar, only the graphs related to the first age group is presented. Fig. 2 shows the average MSE of a male and a female for different dimensions. According to this figure, the MSE difference between the two genders is approximately independent of dimension. Hence, *m* = 5 is selected for the following analysis.

**Fig. 2:**
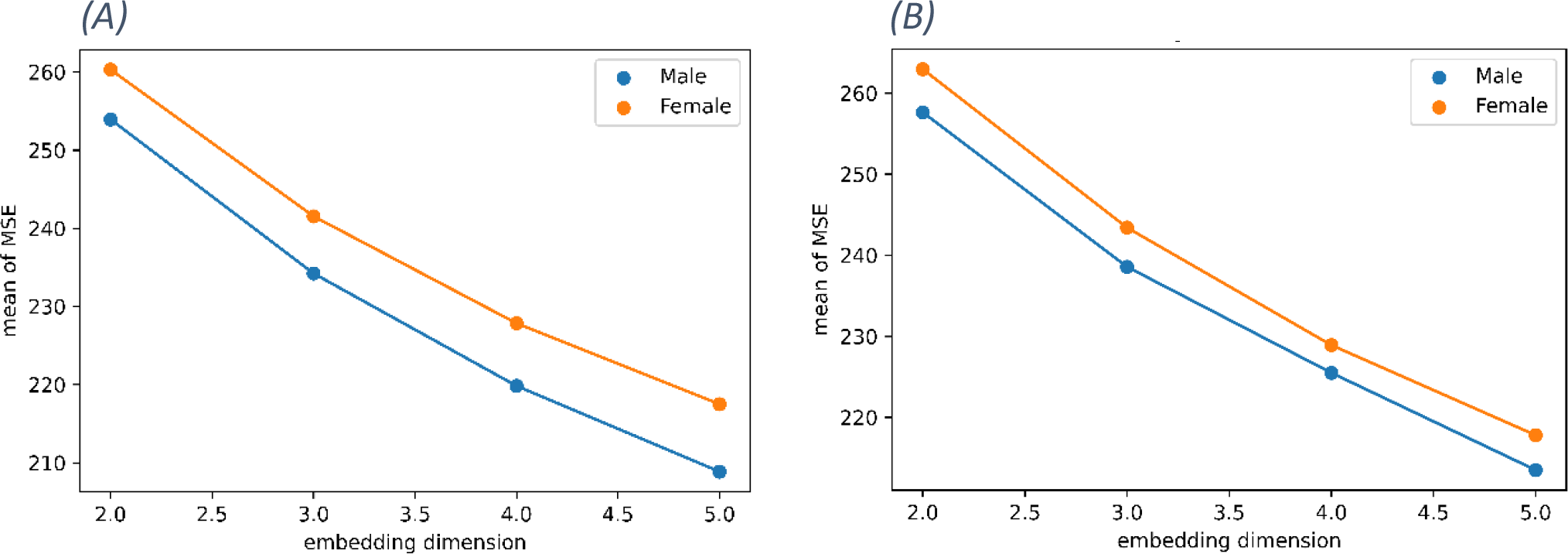
Variations of the mean of MSE with respect to the embedding dimension for (A) all sensors and (B) one randomly selected sensor.

Moreover, to investigate the effect of scale factor(*τ*), one male and one female subject were randomly selected from each age group. For each participant, the sample entropy of all magnetometer sensors up to scale 125 was calculated and averaged over all sensors. Fig. 3 shows the mean of sample entropy in terms of scale for a 23-year-old female and male subject for different embedding dimensions. *τ* = 125 was the largest scale at which the sample entropy of coarse-grained time series becomes finite for all of the sensors. Therefore, the maximum dimension is considered to be 125.

**Fig. 3:**
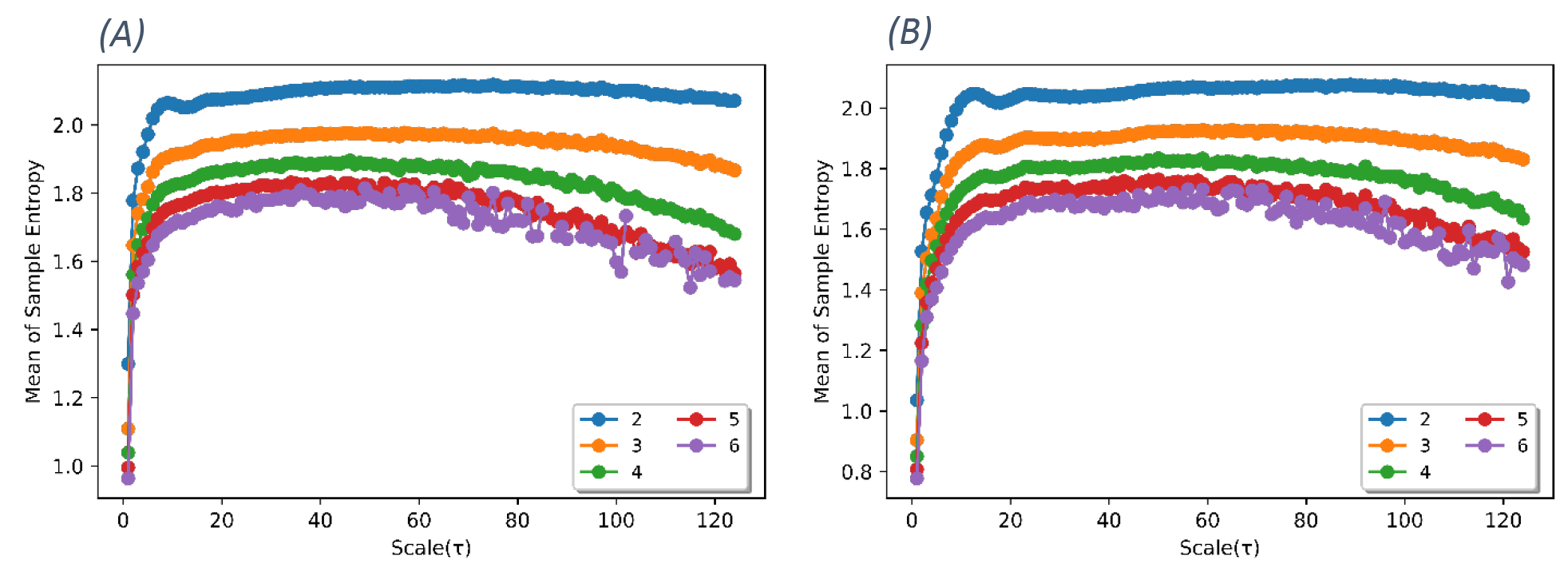
mean sample entropy in terms of scale (up to 125) for (A) a 23-year-old female and (B) a 23-year-old male subject. Different colors are related to different embedding dimensions.

### 3.2. Relationship between age and MSE

Each sensor’s time series was pre-processed as mentioned before. Using the analysis of variance, the result of sensor selection is presented in Fig. 4. Selected sensors were marked with white hollow circles, and other sensors were marked with black dots. Then the mean complexity of chosen sensors was calculated.

**Fig. 4:**
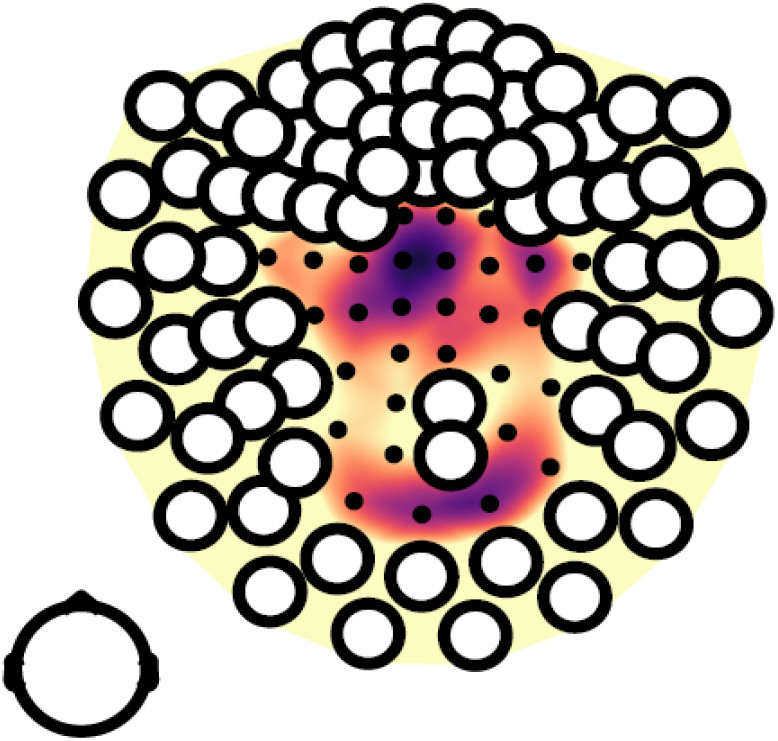
Selected sensors. White circles and black dots represent the selected and non-selected sensors, respectively.

Fig. 5 shows the scatter plot of the mean complexity versus age. As shown in this figure, the complexity is negatively correlated with age (*pearson correlation* = −0.34). The blue line shows the regression line obtained using the least-squares method (P − value < 0.001, slope =−0.227 ± 0.025). The blue halo also indicates the 95% confidence interval. The mean complexity of male and female subjects is presented in Fig. 6. As shown in this figure, males and females display no significant differences in the mean complexity across all age groups. This figure also illustrates that the rsMEG complexity starts declining from the age ∼48 years in both gender groups.

**Fig. 5:**
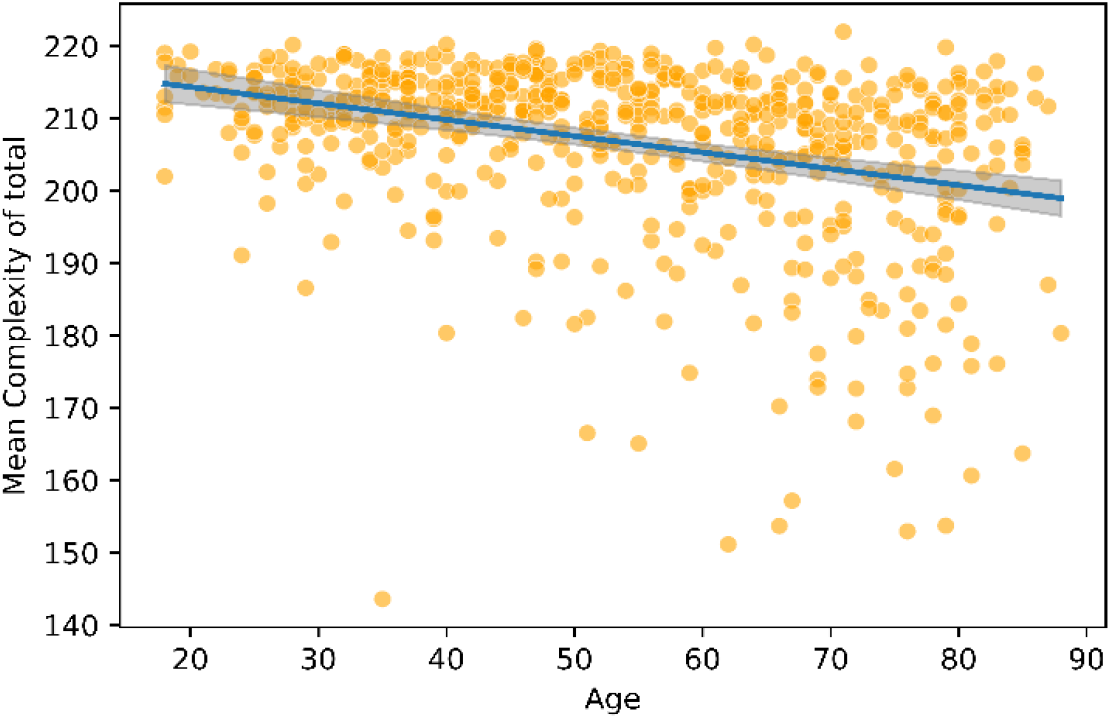
Scatter plot of the mean complexity versus age. The blue line is the least square regression line. The blue halo indicates the 95% confidence interval.

**Fig. 6:**
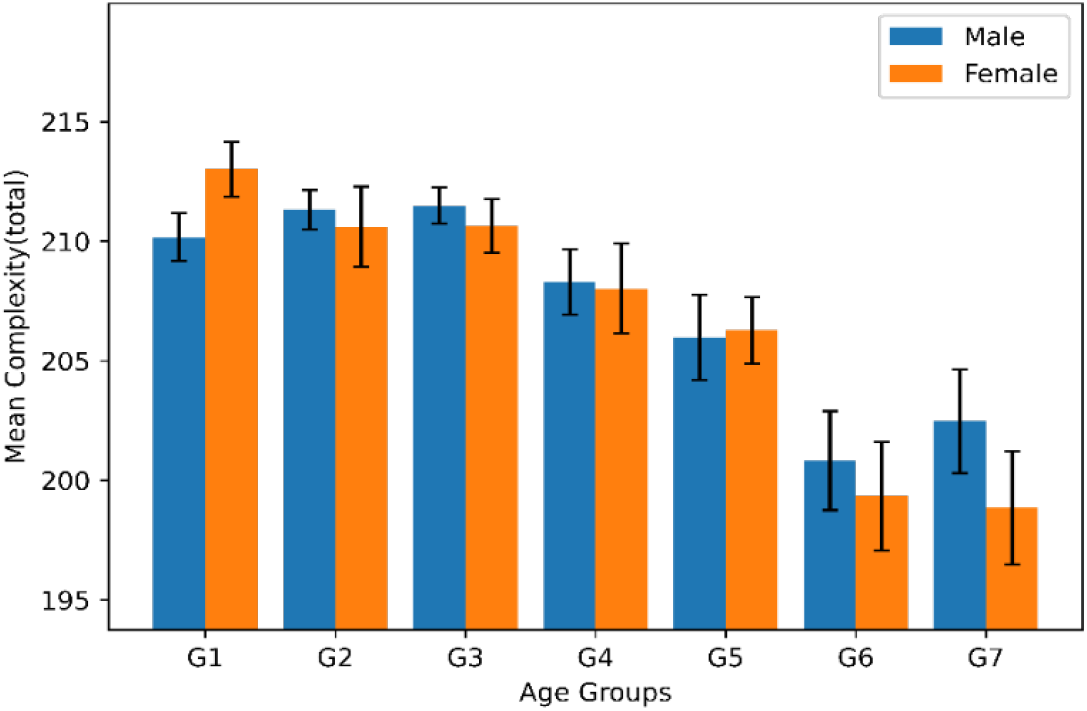
Mean complexity of male and females subjects. Error bars denote the standard error of the means.

### 3.3. Correlation of complexity difference between right-left hemispheres with age

In order to investigate the correlation between the complexity difference of the right-left brain hemispheres and the age, only symmetric channels were selected from the significant sensors identified in Fig. 4. Then, the mean complexity of the symmetric sensors in each hemisphere was calculated. The scatter plot of the complexity difference of the hemispheres is shown in Fig. 7. According to this figure, the difference of complexity between the hemispheres in resting-state healthy participants is positively correlated with age (*pearson correlation* = +0.24). In this chart, the blue line represents the fitting line obtained from the linear regression by the least-squares method (*P* – *value* < 0.001, *slope* = 0.046 ± 0.008). The blue halo indicates the 95% confidence interval.

**Fig. 7:**
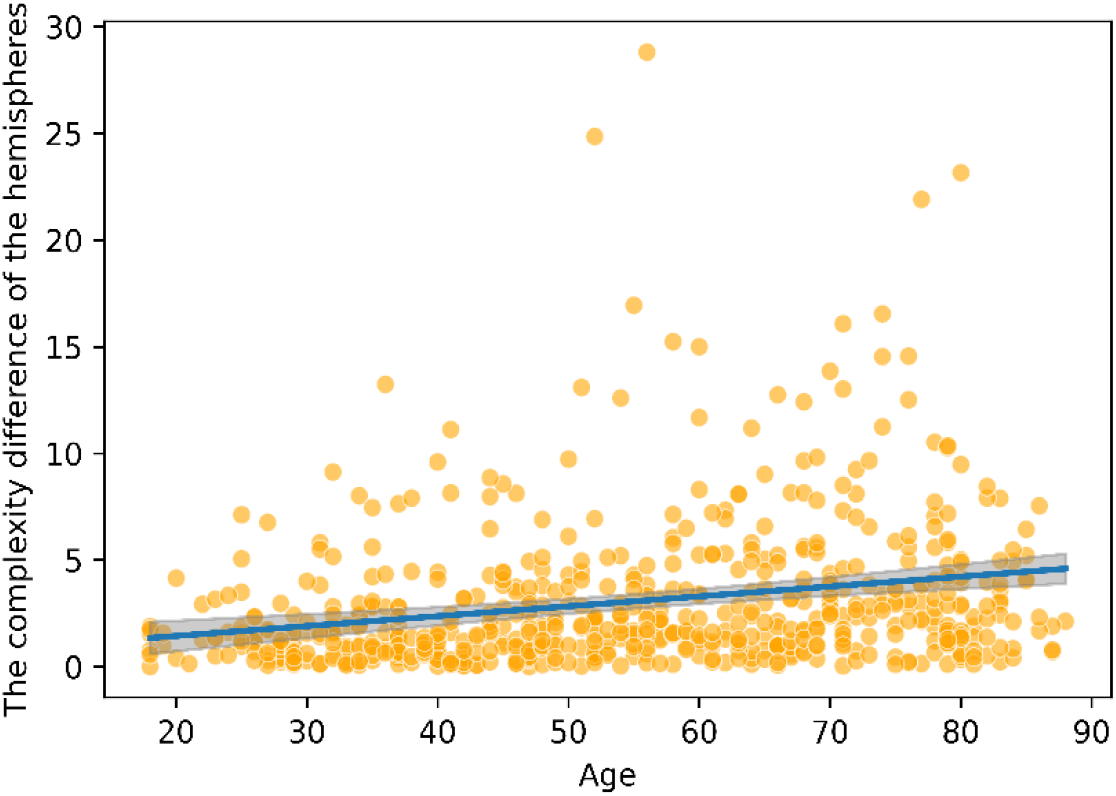
Variations of the complexity difference between the left and the right hemispheres with age.

Fig. 8 shows the effect of gender on the complexity difference between the right-left hemispheres for different age groups. The complexity was averaged only on selected sensors (for reference: See Fig. 4). According to this graph, except for the G4 and G6 groups, there was no significant mean complexity between hemispheres, across males and female subjects.

**Fig. 8:**
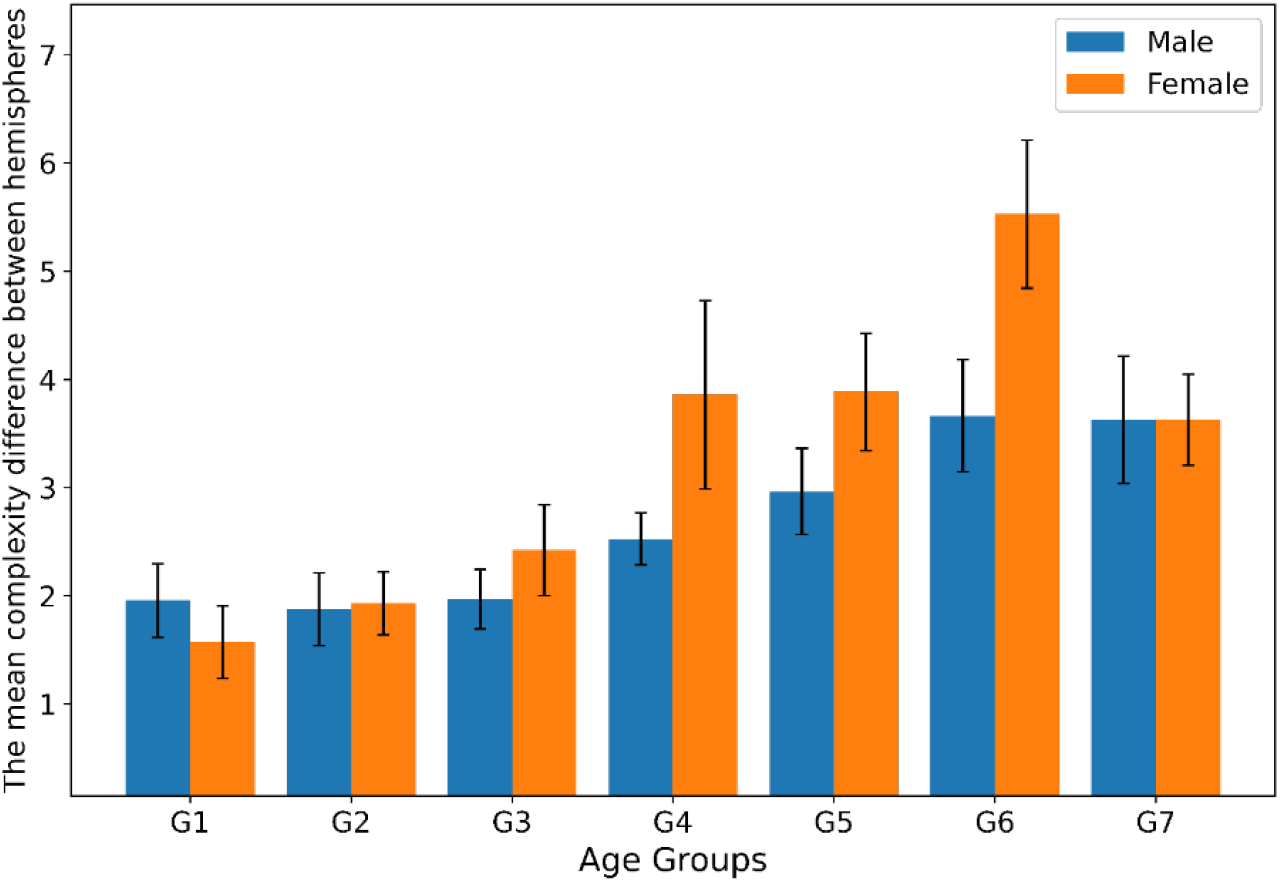
Mean complexity difference between the left and the right hemispheres in 7 age groups under study.

### 3.4. The relationship between complexity and fluid intelligence

The effect of age on fluid intelligence has been previously studied [26, 27]. As shown in Fig. 9, fluid intelligence decreases with age. According to the statistical analysis, the *P*-value is less than 0.001, and the correlation coefficient is -0.66. In this study, different age groups were investigated for the effect of gender on fluid intelligence. As shown in Figure 10, there was no significant difference between males and females in all age groups. Additionally, fluid intelligence begins to decline from the third age group.

**Fig. 9:**
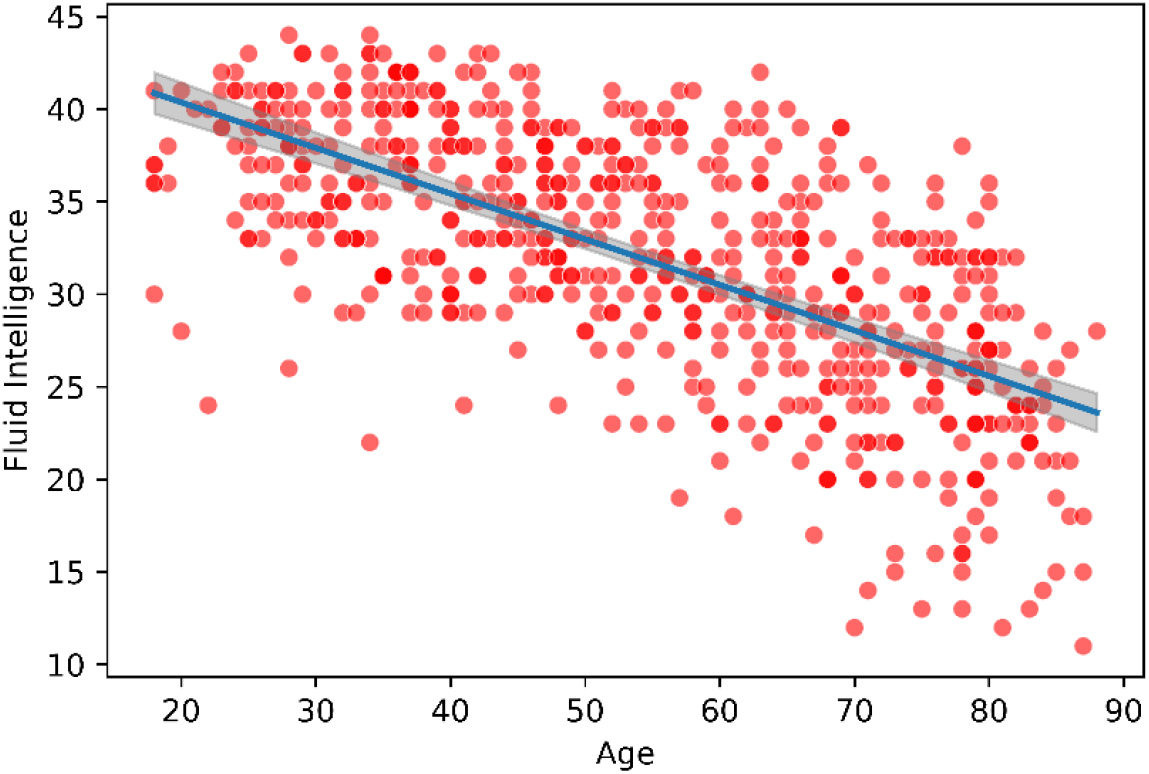
Variations of the fluid intelligence with age.

**Fig. 10:**
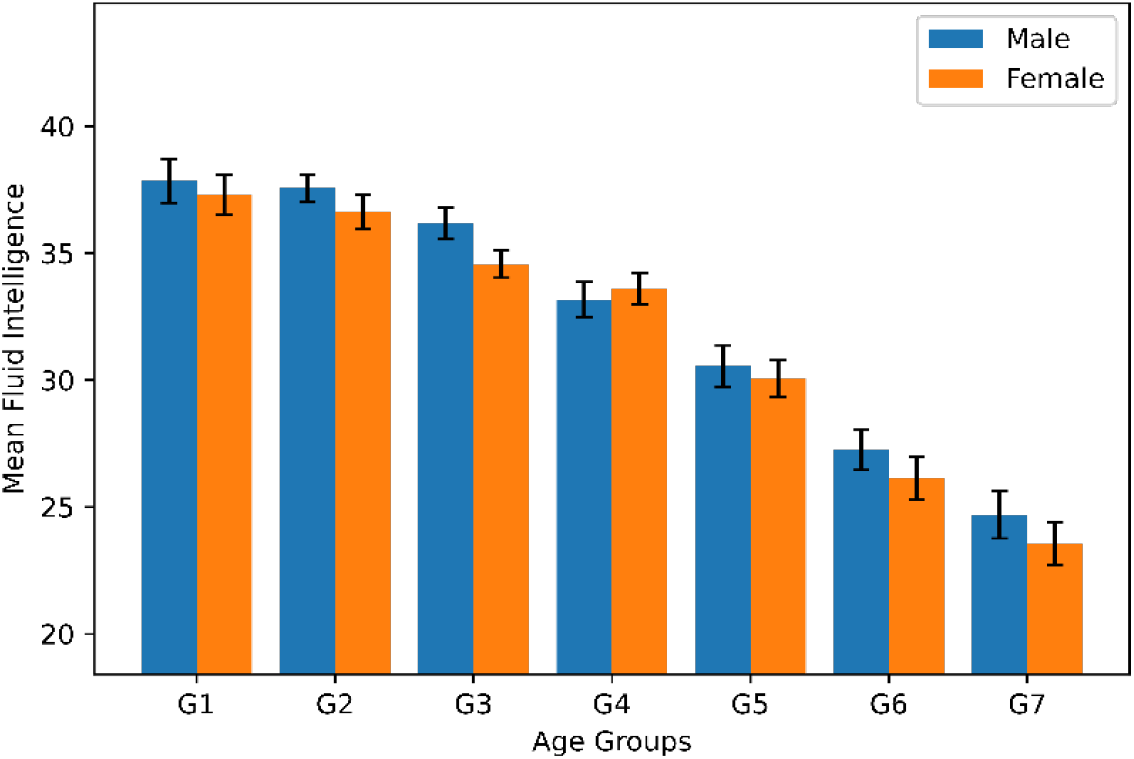
Mean Fluid Intelligence in each age group. Error bars denote the standard error of means.

In Fig.11, the relationship between fluid intelligence and MSE of the rsMEG signal is investigated. The results show a positive correlation between fluid intelligence and complexity (*pearson correlation* = +0.28). Due to the negative correlation between age and complexity, this result was predictable. In Fig. 11, the regression line and the 95% confidence interval are also shown (P − *value* < 0.001, slope = 0.5 ± 0.07).

**Fig. 11:**
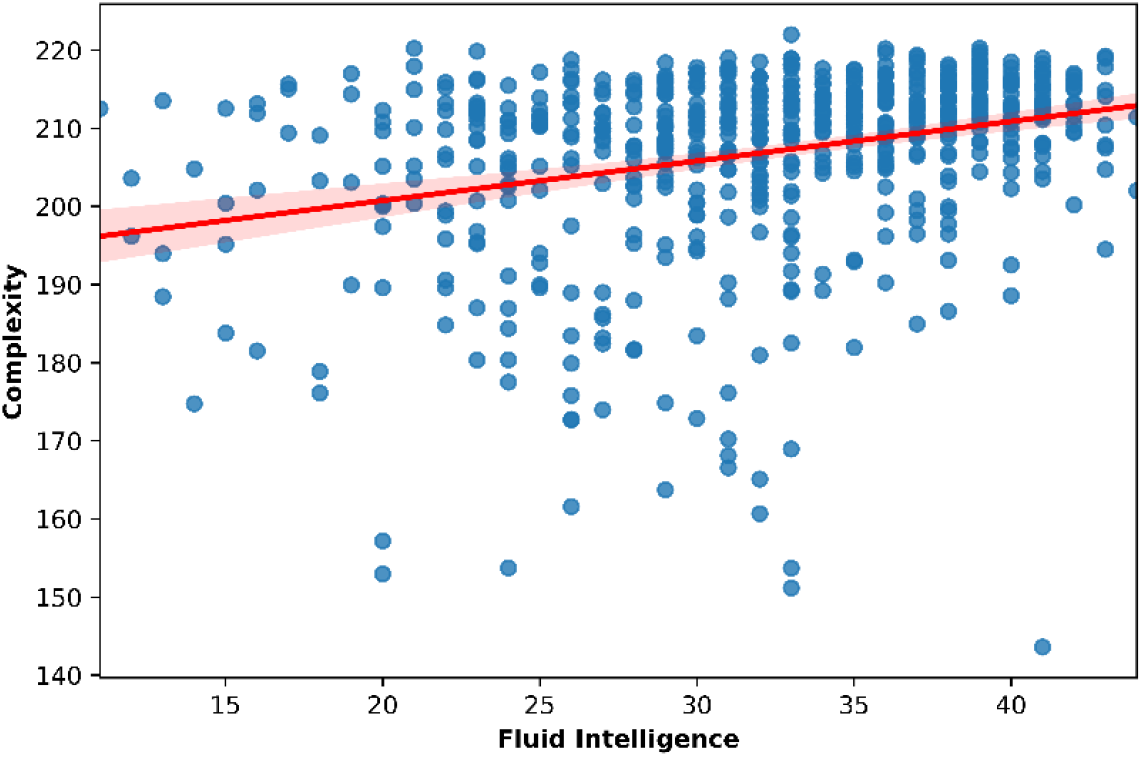
Variations of the complexity with fluid intelligence

### 3.5. Relationship between age and relative power

In this section, the relation between age and MEG relative power is investigated. The relative power was averaged over all sensors and the participants in each age group. This quantity is computed for six bands including alpha, beta, delta, theta, low gamma and high gamma. In Fig. 12, the relative power relationship with age and the differences between males and females were examined. It can be seen that the relative power of the alpha band is higher for the males compared to females in most age groups. However, in the low and high gamma bands, the relative power of females is significantly larger than that of males.

**Fig. 12:**
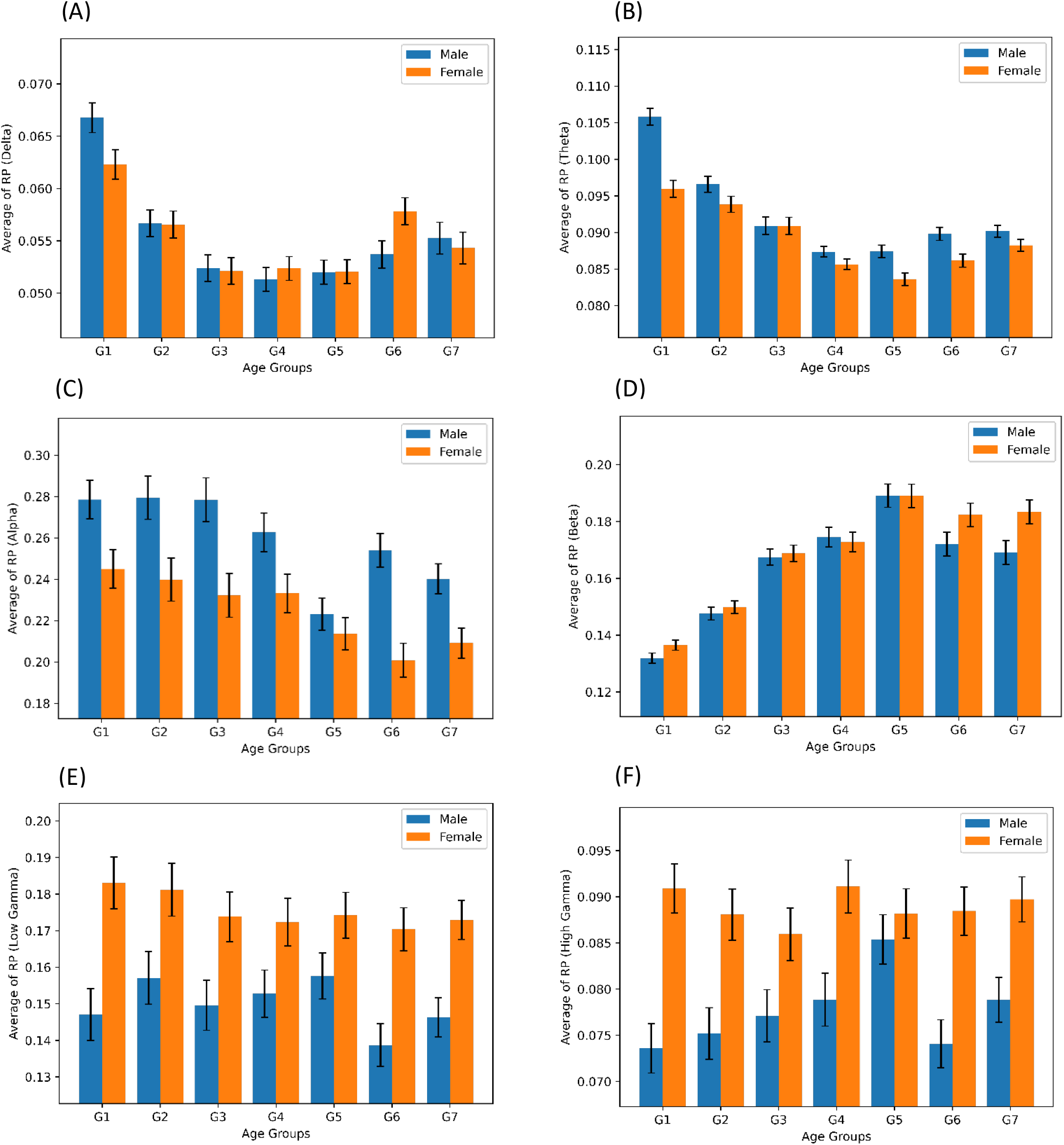
Average relative power in each age group for (A) delta, (B) theta, (C) alpha, (D) beta, (E) low gamma and (F) high gamma bands. Error bars denote the standard error of means.

## 4. Discussion

This study investigated the complexity of rsMEG and Fluid Intelligence in healthy aging. The complexity of the rsMEG signal was characterized using multiscale entropy. Compared to many complexity measures, multiscale entropy is more appropriate due to its robustness to noisy and short time series and the ability to identify complex fluctuation patterns in different temporal scales [14, 23]. Our results showed that the complexity of rsMEG decreases during the aging process. This decrease in complexity of rsMEG is related to various structural and functional changes in the brain across the life span. Some age-related structural changes include a reduction in volume, density, and thickness of gray matter and white matter [28-31]. These structural changes during aging lead to functional changes in the brain. Investigations on functional connectivity of the brain shows that brain activity changes in different areas during aging [29, 32].

Our results are consistent with Matthew King-Hang Ma et al.’s study [33], in which the authors have shown the largest Lyapunov exponent of eye-closed resting-state EEG decreases across the life span. In addition, they showed that complexity quantified by another measure called Lempel-Ziv complexity increases with age. Such a difference is due to the difference in the theoretical definition of complexity measures. Consequently, the theoretical background of different complexity measures should be considered. In another study, which used the Higuchi fractal dimension to quantify the complexity of eye-opened rsEEG, a concave U-shape relationship was obtained between the complexity and age [34]. In addition that the different results may be due to the eye condition (opened and closed eyes); it is also possible that this discrepancy is due to the inappropriate choice of the brain complexity measure since the Higuchi dimension is a measure of monofractal analysis, and affected by the standard deviation of the signal. In comparison, the criteria related to multifractal analysis are better for describing the dynamics of brain signals [35]. Besides complexity reduction in aging, studies have shown that complexity of rsMEG decreases in many neurodegenerative disorders, such as Parkinson’s and Alzheimer’s, and some mental illnesses, such as schizophrenia [36-38].

We showed that the asymmetry in complexity positively correlates with age. Many researchers have revealed structural and functional asymmetries between the left-right hemispheres of the brain [39-42]. One reason for this is the “Yakovlevian torque” phenomenon, in which the right side of the brain tends to deviate slightly forward, and the left side of the brain shifts backward [43, 44]. As a result, the difference in complexity between left-right brain hemispheres most likely arises from these asymmetries. Furthermore, we demonstrated that fluid intelligence and complexity have a positive correlation. Earlier research on fMRI data showed that more entropy leads to more intelligence (not specifically fluid intelligence). It was shown that the sample entropy of fMRI in a resting state carries information about people’s intellectual ability and is consistent with the results obtained in our study [45].

Finally, we observed that the relative powers of the delta, theta and alpha bands decrease with age in the eye-closed resting state. Our results were consistent with previous studies [46, 47]. In addition, our findings show that in the eye-closed resting state, the relative power of low and high gamma rhythms in all age groups for females is higher than the corresponding relative power in males.

## Notes

https://www.cam-can.org/index.php?content=dataset

### Competing Interest Statement

The authors have declared no competing interest.

